# Extended pre-training of histopathology foundation models uncovers co-existing breast cancer archetypes characterized by RNA splicing or TGF-β dysregulation

**DOI:** 10.1101/2025.03.27.645192

**Authors:** Lisa Fournier, Garance Haefliger, Albin Vernhes, Vincent Jung, Igor Letovanec, Pascal Frossard, Cédric Vincent-Cuaz, Raphaëlle Luisier

## Abstract

In recent years, histopathology foundation models (hFM) have rapidly advanced in size and complexity, achieving excellent performance in tasks such as cancer diagnosis and biomarker discovery. Here, we reveal novel capabilities of these models by specializing hFMs, originally trained on diverse tissue types, specifically to invasive tumor tissue. It enables unprecedented discrimination of visually similar yet molecularly distinct tumor regions, that were previously indistinguishable by baseline models, which eventually leads to uncovering new biological insights into breast cancer. Our contributions are threefold. First, to the best of our knowledge, this is the first study to systematically evaluate the biological concepts encoded within hFM representations across multiple scales. Second, we explore extended pre-training to identify optimal conditions that enhance the model’s ability to encode richer, tumor tissue-specific biological concepts. We show that this refinement strategy transforms generalist models into a specialist one capable of resolving subtle, recurrent tumor regions with distinct morphological and molecular identities, called tumor archetypes. Finally, leveraging this specialized model, we uncover two dominant tumor archetypes in invasive breast cancer characterized by distinct aberrant gene expression signatures, notably RNA metabolism dysregulation and TGF-β signaling. Strikingly, these archetypes coexist within the same tumors as spatially distinct regions with varying densities and patterns, and are recurrent across patients, highlighting their universality and potential clinical relevance for patient stratification. Altogether our study demonstrates how extended pre-training of state-of-the-art hFM with specific tumor tissues can unlock rich molecular and morphological information encoded in H&E images. By providing a more accessible approach to investigating tumor heterogeneity, this work opens new avenues for precision oncology, using routine histopathology slides and low computational resources.

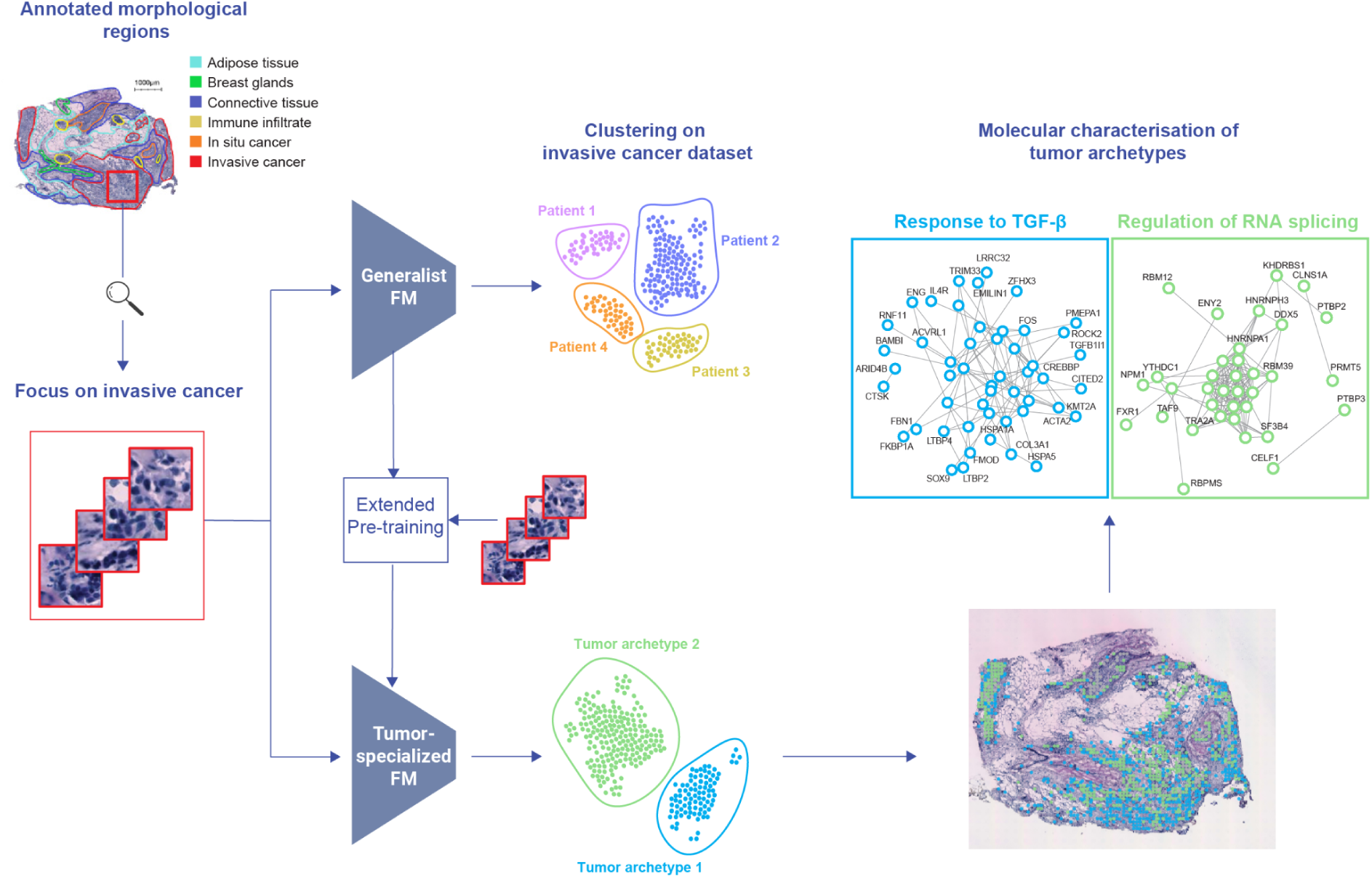
Graphical abstract.

## Introduction

Intratumor heterogeneity (ITH) is a hallmark of cancer^1^. Accurately quantifying and characterizing ITH is essential for improving patient stratification and predicting therapeutic outcomes. ITH has been extensively studied at the molecular level, particularly through single-cell RNA sequencing (scRNA-seq)^2,3^, revealing the coexistence of distinct tumor subpopulations within the same tumor, each defined by unique molecular identities linked to tumor invasion and therapy resistance^4–6^. However, how these distinct molecular programs spatially co-exist within tumors, forming regions defined by unique molecular profiles, tissue organization, and composition, often referred to as “archetypes”, remains poorly understood. Recent advances in spatial transcriptomics (ST) and multi-omics technologies have begun to unravel the complexity of tumor ecosystems by enabling the spatial mapping of molecular programs within tumors^7^. These studies have highlighted how tumor cell populations co-exist within distinct immune microenvironments, underscoring the critical role of spatial context in therapeutic outcomes^8,9^. However, most work to date has focused on tumor-immune interactions, and does not solve the question of whether, and how, tumor cell subpopulations themselves organize into distinct tumor archetypes with potential clinical relevance. While molecular subtypes already guide preclinical models and treatment decisions in many cancers ^9^, a scalable method to map the abundance and spatial organization of tumor archetypes could be transformative for designing more rational combination therapies.

ITH extends beyond molecular variation to include morphological diversity in cell shape and organization, playing a critical role in cancer studies ^10–14^. Importantly, many of these features are reflected in hematoxylin and eosin (H&E)-stained histopathology images, which remain a cornerstone of diagnostic pathology thanks to their rich morphological detail, affordability, and ease of use, making them a powerful and scalable modality compared to more complex techniques like spatial transcriptomics (ST). For example, breast cancer grading is based on multiple morphological criteria, including the degree of nuclear atypia, architectural patterns, and mitotic activity ^15^. The full potential of histopathology to resolve *molecular* ITH, particularly in terms of underlying gene expression programs, remains however largely untapped, often due to the perception that it lacks the molecular resolution provided by techniques like ST ^16^. This perception is reinforced by the fact that cancers with profoundly different molecular programs, and consequently distinct clinical behaviors and therapy responses, can appear morphologically indistinguishable to the human observer, not because the visual information is absent, but because the human brain is inherently limited in processing the full complexity captured in histological slides, often focusing only on diagnostically salient features^9^. However, recent breakthroughs in artificial intelligence (AI), particularly large-scale self-supervised models designed for histopathology, known as histopathology foundation models (hFMs), are nowchallenging this view. These models demonstrate the ability to extract molecular-level information (gene expression profiles and genetic mutations) directly from routine H&E images ^17–22^. These results are consistent with growing evidence that morphological features, such as cell size, nuclear morphology, granularity, and spatial organization, are closely linked to gene expression programs ^23,24^. Collectively, these studies demonstrate that H&E images contain rich biological information, including molecular signals essential for resolving tumor heterogeneity, which can actually be retrieved using hFMs.

While hFMs have shown remarkable potential across tasks, from segmentation^25^ and survival prediction^26^ to tumor characterization ^27–31^, critical gaps remain in understanding what biological concepts are truly captured across biological scales. Fully harnessing their potential requires uncovering how semantic information, defined here as biologically meaningful representations of tissue architecture, cellular composition, and molecular programs, is encoded within these models. Gaining this understanding is crucial for identifying how modifications to model architecture or learning processes impact the type of information captured. This knowledge enables targeted adaptations to ensure models perform in ways that are biologically meaningful and relevant to the task.

In this study, we push the boundaries of hFMs to demonstrate their untapped potential in uncovering molecularly distinct tumor regions invisible to human experts and uncover new biological insights into the molecular composition of invasive breast tumors. First, by systematically comparing six state-of-the-art pathology models, SimCLR ^32^, CTransPath ^33^, Virchow2 ^34^, Prov-GigaPath^18^, UNI ^17^, and its updated version UNI2-h, we provide a comprehensive evaluation of their representational capacity in capturing biological relevant concepts. Next, we show that extended pretraining, which consists in continuing the pretraining phase with additional unlabeled invasive tumor tissue data, enhances the model’s ability to encode richer semantic representations. This refinement transforms generalist models into more specialist ones capable of resolving subtle, recurrent tissue archetypes that are morphologically very similar and shared across patients, but that exhibit distinct molecular identities, as validated by ST. In contrast, conventional hFMs fail at this task, as they tend to identify archetypes with overlapping molecular profiles, and dominated by patient-specific effects, rather than capturing universal archetypes recurring across patients. Crucially, analyses using our specialized hFM uncover that two dominant and clinically relevant molecular pathways in invasive breast cancer, TGF-β signaling and alternative RNA splicing dysregulation, are associated with distinct tumor archetypes. Our study demonstrates that these archetypes co-exist within the same tumor, vary in abundance, and are specific to invasive cancer, a previously unrecognized feature of tumor heterogeneity. Our findings together highlight how AI-driven histopathology can unlock rich molecular and morphological information encoded in H&E images offering new avenues for patient stratification and therapeutic targeting, all from widely accessible histopathology slides and low computational resources.

## Results

### Assessing capacity of histopathology FM to encode relevant biological information

In recent years, histopathology foundation models (hFMs) have become powerful tools, enabling significant advances in automating tasks, including segmentation^25^, whole slide image (WSI) classification, and survival prediction^26^. By leveraging vast amounts of H&E data, these models learn to extract, without relying on predefined labels, rich vector representations from histopathology images related to complex patterns. These models vary widely in architecture and scale, hence potentially in the complexity of their learned representations, with recent versions like Prov-GigaPath exceeding one billion parameters. Benchmarking efforts have primarily focused on standard tasks such as segmentation or whole-slide image (WSI) classification^17,18,32–34^. However, these evaluations are generally performed with the aim of maximizing performance, without a strong focus on studying the learned representations and their biological relevance. Although hFMs learn high-dimensional representations that can capture diverse discriminant features, these may not always align with meaningful biological concepts or be relevant to the studied specific tissue types. Thus we first conducted an analysis to characterize and quantify the semantic information captured by six widely used hFMs (UNI, UNI2-h, Virchow2, Prov-GigaPath, CTransPath, SimCLR; **Supplementary Table 1**), using a well-known publicly available breast cancer dataset^35^ (**Fig. 1a**). This data-set includes H&E-stained image data, expert annotations of six major tissue domains (breast glands, connective tissue, adipose tissue, immune infiltrate, invasive cancer, in situ cancer), and spatial transcriptomic data for 6 patients. After image pre-processing, we extracted patch-level vector representations from each model, as well as segmentation-based handcrafted features (**Methods**), to enable downstream interpretability analyses focused on understanding feature relevance across several biological scales (**Fig. 1**; **Supplementary Tables 1,2**).

**Figure 1.**
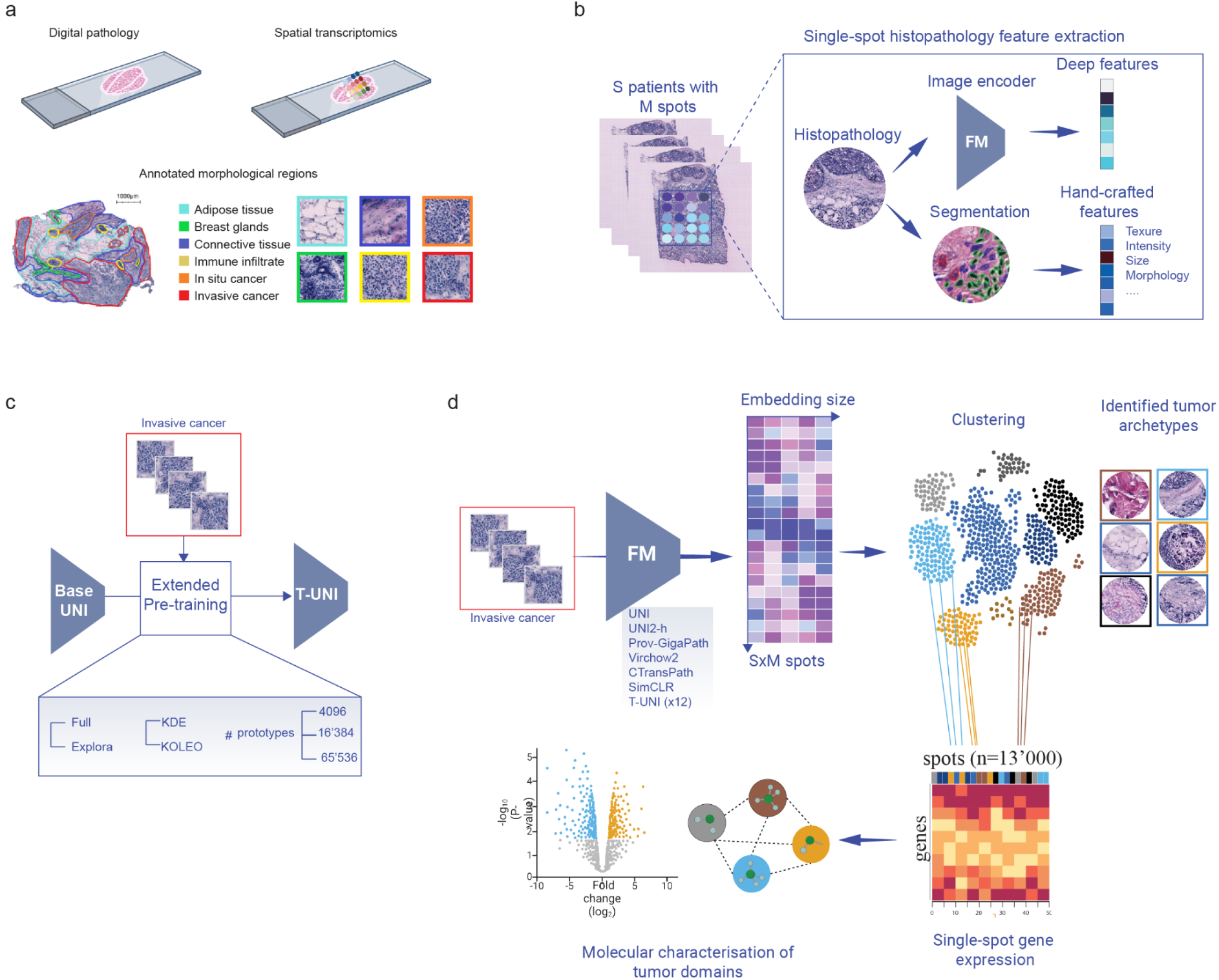
| Overview of the strategy. **(a)** Schema of the comprehensive, publicly available dataset of HER2-positive breast tumors used in this study^35^, comprising H&E images, spatial transcriptomics data, and expert pathological annotations categorizing tissues into six distinct types: adipose tissue, breast glands, connective tissue, immune infiltrate, in situ cancer, and invasive cancer, from eight patients. **(b)** From the H&E image patches, deep features are extracted using histopathology foundation models. In parallel, nuclei segmentation and classification is performed using CellViT^25^, and patch-level handcrafted features are computed to reflect nuclei composition, texture, color and morphology as well as extracellular and whole patch colors and textures. **(c)** Extended pre-training of UNI was conducted using invasive cancer patches only, leading to the tumor-expert T-UNI models. These models differ in their extended pre-training strategies, which include either ExPloRa or full retraining. Additionally, they vary in their regularization losses, utilizing either KDE or KoLeo, and in the number of prototypes, which can be set at 4,096, 16,384, or 65,536. (**d)** Invasive cancer patches vector representations are extracted from the baseline hFMs and T-UNI models. Clustering using UMAP^36^-kmeans is performed to identify tumor archetypes in the image embedding spaces, which are then molecularly characterized by integrating the ST data. Tumor archetypes are described through biological pathways enrichment analysis after marker genes extraction through differential gene expression analysis.

Visual inspection of the Uniform Manifold Approximation and Projection (UMAP) embeddings^36^ of patch-level representations revealed that major tissue components, such as adipose tissue, cancer regions, and immune cell infiltrates, naturally form clusters (**Fig. 2a**). This indicates that all models, regardless of their size, capture global patch properties which are key for differentiating these tissue types, and are actually related to color and nuclear density. However, some models produce clusters with greater biological coherence, grouping similar tissue types more consistently, which suggests better alignment between their learned representations and meaningful biological distinctions. To systematically test this, we evaluated each model’s ability to cluster similar tissue types by computing Adjusted Rand Index (ARI) scores against ground truth annotations, and further examined their correlation with the complexity of each model’s representation space, quantified by Shannon Entropy (SE). SE was computed from the variance ratios explained by the principal components of each hFM’s representations, modeled as a probability distribution (**Methods**). A high SE indicates that information is more evenly spread across the *N* dimensions of the representation space rather than being concentrated in a few. We hypothesized that models utilizing the full capacity of the representation space would exhibit increased semantic richness. As shown in **Fig. 2b**, SE varies significantly across models and correlates to some extent with the representation space dimensionality, as well as model size, with UNI2-h (0.6 billion parameters) and Prov-GigaPath (1.1 billion parameters) exhibiting the highest SE.

**Figure 2.**
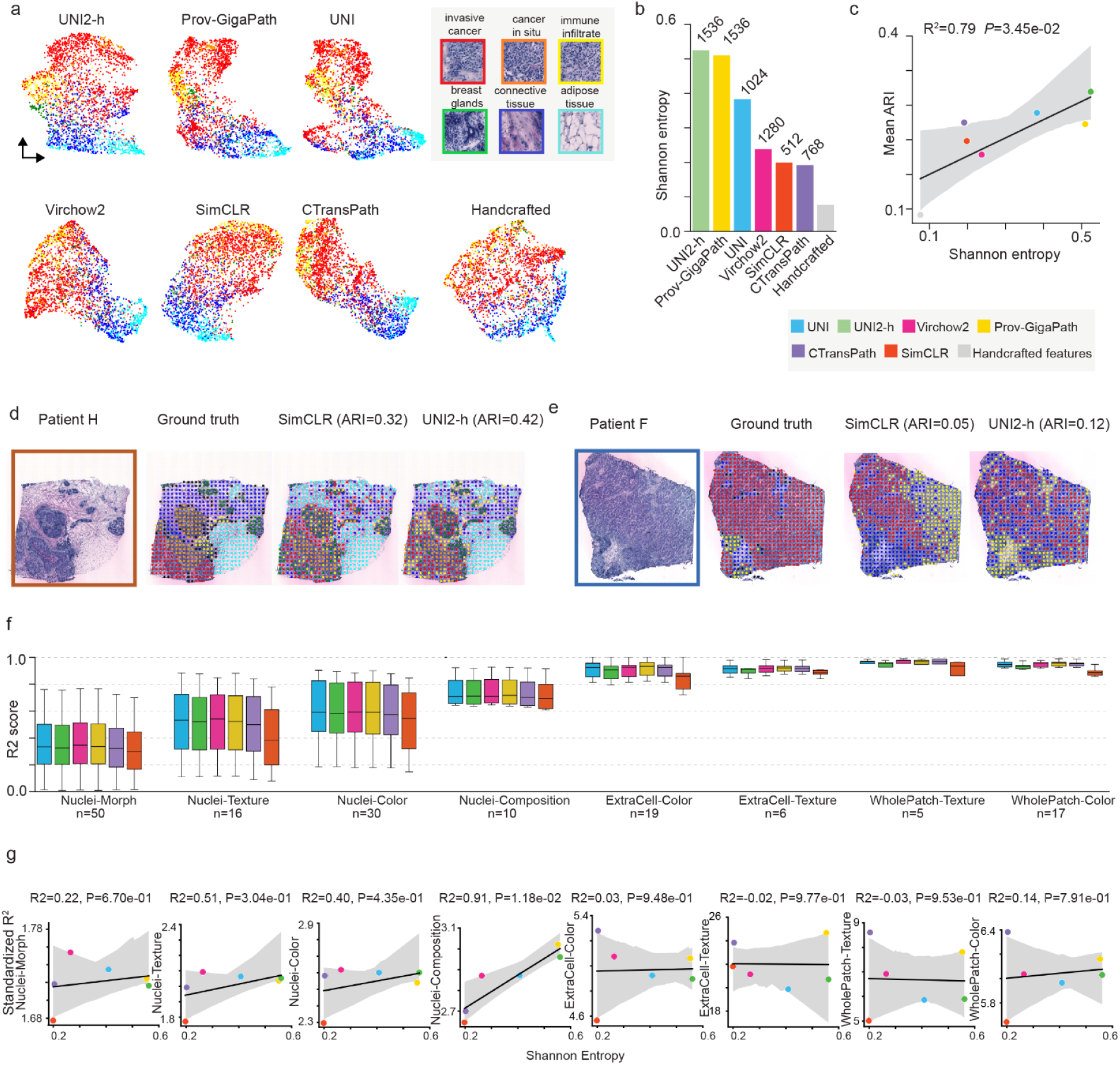
| Assessing semantic richness in histopathology representations with entropy. **(a)** UMAP visualizations^36^ comparing the patches embeddings (*n*_*patc*ℎ*es*_ = 103’023) extracted from the 6 baseline hFMs and the patches handcrafted features, colored by tissue type as annotated by the pathologist. (**b)** Bar graphs representing the Shannon Entropy (SE) associated with the representations provided by each hFM, as well as handcrafted features. Embedding size is indicated at the top for each hFM. **(c)** Scatter plot showing the correlation between SE and mean Adjusted Rand Index (ARI) score across patients for the different hFMs and handcrafted features. ARI measures tissue type discrimination. R^2^ indicates the Pearson correlation coefficient and P the associated p-value. The line marks the linear regression fit with the 95% confidence interval as error band. Clustering is performed using UMAP-kmeans, with UMAP parameters optimized for the best ARI score. **(d)** Whole slide image (WSI) with ground truth spots annotations and spots clustering (UMAP k-means, k=6) based on SimCLR and UNI2 patches embeddings for patient H. **(e)** Same as **d** for patient F (UMAP k-means, k=3). **(f)** Boxplot showing the distribution of R² scores from linear regression models predicting handcrafted features based on embeddings from different hFM patches. Results are grouped by feature categories: nuclear morphology, nuclear texture, nuclear color, nuclear composition, extracellular color, extracellular texture, and combined color and texture of the entire patch. Number of features is indicated under each plot. Boxplots display the five number summary of median, lower and upper quartiles, minimum and maximum values. (**g**) Scatter plot of the correlation between each feature category standardized R² and SE for the different hFMs. The standardized R² for a feature category is defined as 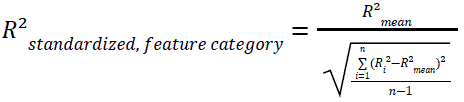 with 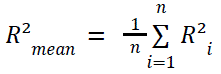, where *n* is the number of features in the feature category and *R*²_*i*_ is the *R*² of the *i-th* feature in the feature category. R^2^ indicates the Pearson correlation coefficient and P the associated p-value. The line represents the linear regression fit, with shaded areas indicating the 95% confidence interval.

Few exceptions are observable, such as UNI and Virchow, suggesting that the former proportionally captures more concepts relevant to distinguish cancerous breast tissues than the latter (**Supplementary Fig. 2d**). Strikingly, we observe that model performance for tissue type clustering is positively correlated with SE, taking into account both non-linear (**Fig. 2c**) and linear (**Supplementary Figs. 2a-d**) relationships between representations. This supports our hypothesis that SE may serve as a reliable indicator of semantic richness. Further analysis of individual ARI scores per patient reveals that most models achieve the highest classification performance on slides with a diverse mix of tissue types, such as in patients H and B (**Fig. 2d**, **Supplementary Fig. 2c**). In contrast, slides dominated by only a few tissue types, such as invasive cancer and small immune cell clusters, pose a greater challenge for all models, as seen in patient F (**Fig. 2e, Supplementary Fig. 2c**), confirming their limitations in capturing fine-grained tissue differences in low-variability samples.

We next evaluated whether these models capture low-level visual features by testing how well their representations linearly predict 153 handcrafted descriptors of color, texture, shape, and cell composition with a focus on nuclear and extracellular matrix characteristics (see **Supplementary Table2** for the full feature list). As shown in **Fig. 2f**, all models except SimCLR perform comparably, consistently showing a stronger ability to linearly encode texture and color-related features than morphological ones. This suggests that pure transformers are inherently more effective at capturing textures and colors than complex morphological features. Moreover, the stronger representation of extracellular features and whole-patch descriptors indicates that these models tend to encode broader tissue context rather than focusing on individual cellular structures. Nevertheless, these models excel at aggregating and denoising cell-related features to accurately infer cell types. This learning pattern aligns with previous studies on natural images, where these models are shown to identify, for instance, texture and color features in the first layers of their architecture, at both local (token-level) and global (patch-level) scales, while the last layers identify high-level discriminant concepts^37,38^. Finally, examining the correlation between SE and each model’s ability to capture different categories of handcrafted features, as summarized using the standardized R² score across all features within each category, we found that higher SE is significantly associated with improved performance in capturing nuclear composition, including the number of nuclei and the density of inflammatory and tumor cells (**Fig. 2g, Supplementary Fig. 2e**). These findings suggest that the enhanced performance in tissue identification observed in models with higher SE is likely driven by their increased capacity to encode semantic information related to nuclear type and density.

Together, these results indicate that SE is a reliable metric for assessing the biological relevance of learned representations, as demonstrated by its significant correlation with the model’s ability to accurately distinguish tissue types based on biologically meaningful differences. Furthermore, the increase in SE, and consequently, semantic richness, is specifically linked to nuclear density and composition.

### Extended pre-training enhances tumor-specific representations while reducing patient-specific bias

Building on our finding that UNI variants and Prov-GigaPath encode a higher variety of biological concepts in their learned representations that can help discriminate tissue, we next focused on the challenge of detecting subtle morphological and compositional differences, hereafter referred to as “archetypes”, within a single tumor compartment. We therefore performed clustering of invasive tumor patches using representations from UNI, UNI2-h, and Prov-GigaPath aiming to detect tumor archetypes with distinct composition and tissue morphology. The analysis identified different numbers of archetypes representing tumor patches across six patients, varying by hFM (**Supplementary Figs. 3b-d**). To ensure biological relevance and generalizability, archetypes should capture patterns consistently observed across multiple patients rather than patient-specific clusters potentially driven by artifacts. However, the archetypes identified using these model representations were poorly distributed across patients, with several individuals predominantly represented by a single cluster, indicating a strong patient effect. These findings suggest that most archetypes identified by these models are driven by technical artifacts or patient-specific factors, or, as in the case of c0 and c2 identified by UNI, may simply represent the same tissue type.

One possible explanation for this poor performance is that resolving archetypes with subtle morphological differences, while avoiding batch effects or patient-specific artifacts during training, requires richer, tumor-specific semantic representations. This underlines the need for models better specialized in tumour tissue encoding which can be obtained by leveraging Extended Pre-learning, also known as self-supervised fine-tuning, which comes down to continuing model training exclusively on that data modality ^39,40^. We therefore tested this method exclusively with UNI, as the training parameters for UNI2-h or Prov-GigaPath were not available at the time of the study. Additionally, UNI requires significantly less computational resources than the other models (e.g., different GPU requirements), making it a natural choice for this first study. Given the lack of established standards for extended pre-training, we systematically explored multiple strategies and hyperparameters (**Methods**). In brief, we compared “full training”, which updates all model weights, with ExPLoRA, a partial weight update approach designed to better preserve prior knowledge. Additionally, we tested two regularizations (KDE^34^ and KoLeo^41,42^) promoting better representation spread. Finally, models were trained using varying numbers for their learned “prototypes”. This resulted in 12 tumor-specialized models, referred to as T-UNI (**Fig. 1c; Supplementary Table 3**). Overall, extended pre-training did not significantly alter the global biological concepts encoded in the learned representation, which remained primarily structured by tissue types, regardless of the training procedure (**Fig. 3a**). However, full training resulted in a clearer separation between invasive tumor tissue patches and other tissue types compared to the UNI baseline and models trained with ExPLoRa. The effect was particularly pronounced with KoLeo regularization, which promotes a more uniform distribution of representations across the embedding space, effectively diffusing them further apart, compared to the smoother impact of KDE (**Fig. 3b**). We further observed that ExPLoRA simultaneously improved tissue clustering performance (increase in ARI scores) and SE, demonstrating that extended pre-training with this method refines representations whilst keeping prior model knowledge of all tissue types (**Fig. 3c**). However, in the full training setup, ARI scores decrease while SE increases, suggesting that, unlike ExPLoRA, which helps mitigate “catastrophic forgetting”, full training causes the model to lose previously encoded biological concepts in learned representations essential for accurate tissue classification (**Fig. 3d, Supplementary Fig. 3a**). We also found that full training with KoLeo was largely insensitive to the number of prototypes, whereas increasing prototype numbers in KDE progressively reduced SE and impaired the model’s ability to distinguish tissue types (**Fig. 3c**). This suggests that excessive smoothing of representations across the model’s embedding space conflicts with the formation of clusters driven by prototypes and is detrimental to models’ ability to learn biologically meaningful distinctions.

**Figure 3.**
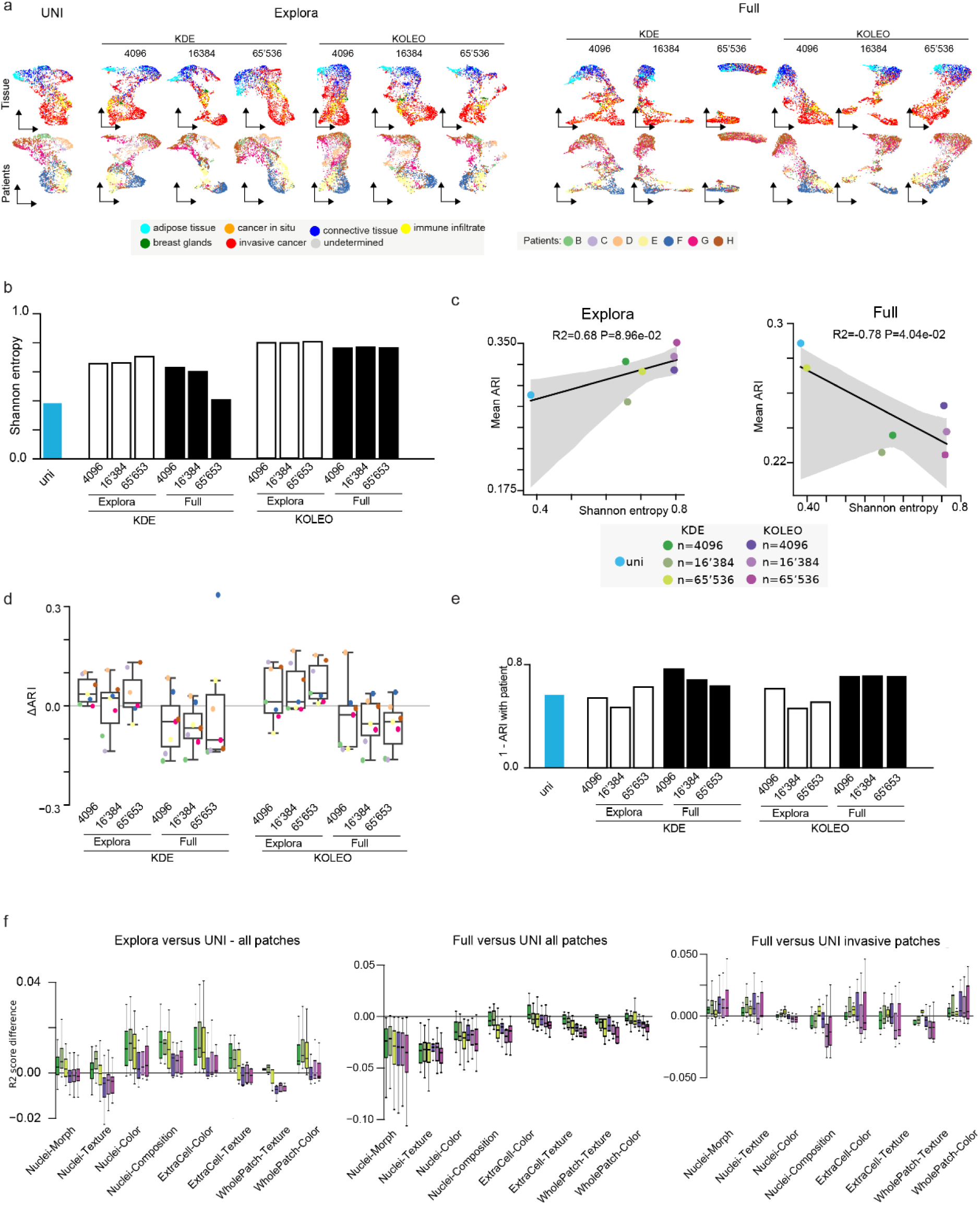
| Extended pre-training exclusively with invasive cancer tissue enhances semantic representation. **(a)** UMAP visualizations^36^ comparing the patches embeddings of UNI and T-UNI models. Top: colored by tissue origin. Bottom: colored by patient origin. **(b)** Bar graphs representing the Shannon Entropy (SE) associated with the representations provided by UNI and T-UNI models. **(c)** Scatter plot of the correlation between SE and mean Adjusted Rand Index (ARI) score across patients for UNI and T-UNI models using ExPloRa method (left) and full retraining method (right). ARI measures tissue type discrimination. T-UNI models trained using KDE loss are in shades of green and models trained using KoLeo are in shades of purples. The different shades represent different prototype numbers (n). Clustering is performed using UMAP-kmeans, with UMAP parameters optimized for the best ARI score, and k set to the number of annotated tissues. **(d)** Boxplots of the distribution of ARI score differences per patient with respect to UNI (baseline) for T-UNI models. ARI per patient shows the model’s ability to discriminate between tissue types from clustering (UMAP k-means) at the patient level. (**e**) Bar graphs of the batch effect mitigation score for UNI and T-UNI models, defined as 1-”*ARI with patient”*, computed using patient labels instead of tissue labels on invasive cancer patches. A high batch effect mitigation score indicates low batch effect. Clustering is computed using UMAP-kmeans. (**f**) Distributions of R² score difference with respect to UNI from linear regression predictions of handcrafted features, grouped by feature category. Linear regression using all patches embeddings for T-UNI models trained with ExPloRa method (left), using all patches embeddings for T-UNI models trained with full retraining method (middle) and using only invasive cancer patches embeddings for T-UNI models trained with full retraining method (right). The box plots indicate the data quartiles with the whiskers extending to the full distribution excluding outliers outside the 1.5 interquartile range while the median is indicated as a horizontal line.

We next aimed to assess whether extended pre-training mitigates the *batch effect* observed with UNI, UNI2-h and Prov-GigaPath, where clusters were dominated by patches from single patients (**Supplementary Figs. 3b-d**). This effect is also evident in UMAP visualizations of patch-level representations from UNI, where patient samples cluster into distinct regions (**Fig. 3a**, *lower*). The goal of extended pre-training on invasive cancer was to enhance the model’s ability to represent this specific tissue type, potentially at the cost of losing relevant information for other tissue types, a trade-off observed with full training but potentially beneficial for identifying tumor archetypes. We argue that reducing the batch effect effect is a strong indicator of improved generalization, as it suggests that the model captures tumor-wide semantic representations rather than patient-specific features, crucial for identifying archetypes shared across patients. We approximated batch effect by defining a *batch effect mitigation score* as 1-ARI, computed using patient labels instead of tissue labels on patches from invasive cancer. As shown in **Fig. 3e**, full training consistently results in lower batch effect, suggesting that these models become agnostic to patient origin.

Having found that extended pre-training increases semantic richness while reducing batch effects, we next sought to determine whether these improvements are linked to specific, interpretable differences, such as enhanced recovery of handcrafted feature representations. Extended trained models with ExPLoRa exhibited an improved capacity to linearly encode specific handcrafted features, particularly those related to cell color, composition, and texture, but at the expense of whole-patch texture representation (**Fig. 3f**). In contrast, models trained with KoLeo systematically showed reduced performance in capturing handcrafted features. However, given that KoLeo enforces a more even distribution of information across the representation space compared to KDE, this lower performance may reflect a shift toward a more nonlinear encoding of these features rather than an actual loss of information. Moreover, we found that full pretraining leads to an overall reduction in the encoding of handcrafted features, which aligns with the observed decline in tissue-type identification performance (**Fig. 3c**). Notably, this effect was reversed when evaluating models on invasive cancer patches alone. This suggests that extended pre-training on invasive tumor tissues exclusively, enhances the model’s ability to capture cancer-specific handcrafted features, while potentially deprioritizing features relevant to other tissue types.

In summary, these analyses indicate that UNI, UNI2-h, and Prov-GigaPath models tend to prioritize global tissue characteristics rather than capturing finer distinctions between tumor regions with distinct molecular profiles. This is compounded by patient-specific effects that impair the detection of subtle archetypes within tumors. Crucially, we demonstrate that while full extended pre-training on invasive cancer patches reduces the model’s ability to distinguish general tissue types (e.g., breast glands, immune cells) and broad handcrafted features, it significantly enhances the model’s capacity to learn tumor-specific biological concepts and effectively mitigates patient-related batch effects. Notably, the consistently superior patient-level ARI scores achieved with KoLeo further support its selection over KDE for this task.

### Full extended pretraining with KoLeo enables the identification of molecularly defined tumor archetypes

Having established that extended pre-training on invasive tumor tissue patches enhances the model’s ability to capture tumor-related biological concepts, we next aimed to test whether these models outperform UNI in identifying universal tumor archetypes, recurrent across patients and most importantly exhibiting subtle morphological differences reflecting distinct molecular identities. We thus performed clustering of tumor patches using representations from all T-UNI models. The analyses identified varying optimal numbers of tumor clusters, ranging from 4 to 9 (**Methods**), with the highest cluster count observed in models trained with ExPLoRA (**Fig. 4a**). Because models trained with ExPLoRA still encode patient-specific information (**Fig. 3e**), it is possible that some clusters derived from these models primarily capture differences between patients rather than reflecting biologically relevant tumor features. To verify this, we next assessed the molecular identities of identified clusters by analyzing their gene expression profiles using spatial transcriptomics data matched to the analyzed H&E tumor patches. First, examining the distances between clusters in gene expression space, we found that most models performed similarly to the UNI baseline (**Fig. 4b**). However, two T-UNI variants, full KoLeo with 16K and 65K prototypes, exhibited increased molecular distances between clusters, suggesting they form more biologically coherent and distinct groups. Further analyzing the average molecular distances between clusters within each patient, as opposed to the overall analysis across all patches and patients (**Fig. 4b**), we found that the T-UNI full KoLeo model with 16K prototypes consistently outperformed all other models in identifying distinct tumor archetypes characterized by defined morphology and molecular identity (**Fig. 4c**). This highlights its superior ability to capture biologically meaningful intratumor archetypes. Based on these results, we selected the T-UNI full KoLeo model with 16K prototypes for further investigation of tumor heterogeneity in breast cancer.

Reproducibility of tissue archetypes across patients is a key indicator of model robustness. Further analysis of cluster distribution revealed that tumor archetypes identified by T-UNI full KoLeo (16K prototypes) are consistently represented across patients, unlike those from UNI, UNI2-h, and Prov-GigaPath (**Supplementary Figs.3b-d**). Notably, two key archetypes, c0 (orange) and c2 (gray), which show the highest morphological and structural similarity based on hierarchical clustering of their H&E learned representations (**Fig. 4d**), were consistently present across all patients (**Fig. 4e**, *upper*). These archetypes also showed distinct spatial organization within tumors, with c0 evenly distributed throughout the tumor mass, while c2 formed localized high-density regions within invasive cancer areas (**Fig. 4e**, *lower*). Hierarchical clustering of their gene expression profiles further confirmed this relationship, showing strong concordance with the image-based representation. In particular, similar to the image results (**Fig. 4f**, *upper*), c0 and c2 consistently exhibited the closest molecular identities in most patients (**Fig. 4f**, *lower*). Given the inherent variability and potential technical noise in clinical data, we hypothesized that robust clusters would exhibit consistent molecular relationships across patients. Specifically, if the hierarchical clustering of gene expression profiles, performed within each patient’s clusters, yields similar structures across patients, it suggests the model captures true biological signals rather than patient-specific artifacts or random variation. To test this, we computed the correlation of hierarchical clustering results based on gene expression data across patients. This analysis showed that models trained with T-UNI full KoLeo (16K and 60K prototypes) not only achieved the strongest batch effect mitigation (**Fig. 3e**) but also produced the most consistent molecular cluster structures across patients compared to other models (**Supplementary Figs. 4a-b**).

These results highlight the superior robustness of KoLeo-based extended pre-training with 16’384 prototypes over baseline models in identifying tumor archetypes with biological significance. The consistent detection of these archetypes across patients, supported by reproducible gene expression profiles, underscores the model’s ability to capture meaningful tumor heterogeneity beyond patient-specific or technical noise.

### Distinct tumor archetypes defined by TGF-β signaling or RNA splicing dysregulation co-exist in invasive breast cancer

Having identified robust tumor archetypes with distinct spatial organization and molecular profiles within tumors, we next sought to evaluate their biological relevance by investigating their molecular identities in greater detail through differential gene expression (DGE) analysis. As an initial validation, we assessed pathway enrichment across the six pathologist-annotated tissue types (e.g. breast gland, connective tissue, cancer in situ) and confirmed that each showed gene signatures consistent with their biological functions, for example, complement activation in immune tissue and vascular regulation in connective tissue (**Supplementary Fig.5a**). This analysis further revealed distinct aberrant molecular programs differentiating *in situ* from invasive cancer. Specifically, DNA repair and chromosome organization pathways were predominantly associated with cancer *in situ*, while invasive cancer was characterized by dysregulation of RNA splicing and activation of TGF-β signaling. Notably, although these molecular signatures are well established in oncology and breast cancer ^5,43–46^, our findings extend beyond previous studies by showing that invasive cancer exhibits exacerbated profiles of these pathways compared to *in situ* disease. This contrasts with initial work with this breast cancer dataset, which primarily focused on the tumor microenvironment rather than tumor-intrinsic molecular programs ^35^.

Having established this baseline, we next focused on the newly identified tumor archetypes. Strikingly, while c0 and c2 are the most closely related both molecularly and morphologically (**Figs. 4d,f**), they display enrichment in biological pathways typically associated with distinct tissue types (**Fig. 5a**). Moreover, each archetype exhibits specific molecular programs, confirming that our approach successfully identified biologically distinct tumor compartments. In particular c0, the most abundant tumor archetype, aligns more closely with invasive cancer, displaying aberrant RNA metabolism (splicing and localization) and energy metabolism remodeling, a well-established hallmark of ER/PR(+) breast cancer^47^ (**Figs. 5a,c**). In contrast, c2, though less abundant yet present across all patients, shows greater resemblance to connective tissue and is strongly associated with TGF-β signaling, a pathway that plays a crucial role in cancer cell proliferation, growth, inflammation, angiogenesis, and metastasis ^45,48^ (**Figs. 5a,b**). Its proximity to connective tissue suggests that c2 may exhibit signatures of invasiveness and epithelial-to-mesenchymal transition (EMT), as connective tissue-related gene programs, including extracellular matrix (ECM) remodeling, collagen production, and vascular regulation, are closely linked to EMT processes^49^. Further analysis of the protein-protein interaction networks for genes differentially expressed in the c0 and c2 clusters confirmed the preferential aberrant activation of TGF-β signaling in c2 (**Fig. 5b**) and RNA splicing pathways in c0 (**Fig. 5c**). The finding that overall trends in differential expression are similar between c0 and c2 is expected given their spatial proximity within the tumor and potential noise in the clustering process. The spatial specificity of these molecular pathways within invasive tumors was further confirmed by visualizing the expression of representative genes, TGFB1I1 for TGF-β signaling and ENY2 for RNA splicing, which displayed distinct spatial activation patterns, reinforcing that these pathways are linked to separate tumor regions (**Fig. 5d**). These findings indicate that tumor archetypes inferred from histopathology do not only recapitulate established cancer subtypes with defined molecular identities but also reveal their co-occurrence in distinct tumor regions within the same patients. Notably, the two dominant tumor archetypes identified using the UNI baseline, while exhibiting similar transcriptional programs and distributions across patients, appear less distinct at the molecular level. In particular, RNA metabolism and TGF-β signaling are equally enriched in both dominant clusters, suggesting that the baseline model lacks the capacity to differentiate tumor subtypes with subtle but biologically meaningful variations (**Supplementary Fig.5c**). This further underscores the value of extended pre-training in uncovering fine-grained morphological differences linked to distinct molecular identities, which may otherwise remain undetected.

**Figure 4.**
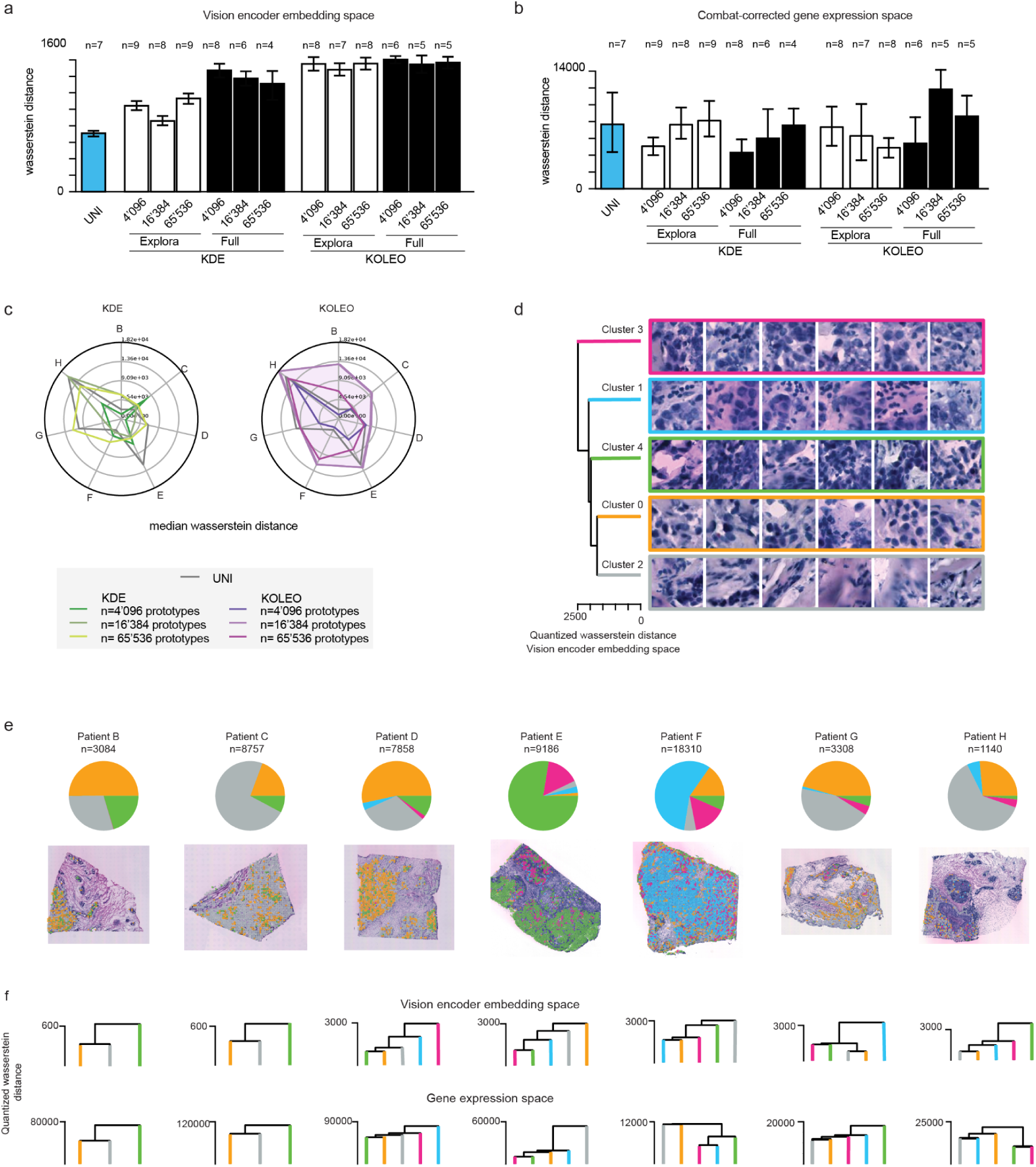
| Full extended pretraining with KoLeo enables the identification of molecularly defined tumor archetypes. (**a**) Bar graphs of the average normalized quantized Wasserstein distances in the UNI and T-UNI embedding spaces between tumor archetype pairs identified from clustering of invasive cancer patches. The 95% confidence interval is shown in black. The number of tumor archetypes identified is indicated at the top for each model. It is selected such as to obtain a maximal silhouette score while maintaining a high batch effect mitigation score (**Methods**). (**b**) Bar graphs depicting the average median quantized Wasserstein distances between archetype pairs identified with UNI and T-UNI models. These pairwise distances were computed for each patient individually in the combat-corrected gene embedding space. The median of these pairwise distances was then calculated for each patient. The bar graphs show the distribution of these median values across all patients, providing the average and the 95% confidence intervals. (**c**) Radar plot showing the median quantized Wasserstein distances between archetype pairs identified with UNI and T-UNI models, computed as in **b**. (**d**) Hierarchical clustering (quantized Wasserstein distance and Ward clustering) of the tumor archetypes identified with the T-UNI model using full retraining, KoLeo loss and 16K prototypes, with representative example patches for the different tumor archetypes. The quantized Wasserstein distance is computed in the T-UNI embedding space. (**e**) Pie chart showing the relative abundance of the 5 tumor archetypes identified with the T-UNI model using full retraining, KoLeo loss and 16K prototypes. Top: Relative abundance of morphological tumor archetypes per patient. Bottom: Invasive cancer patches colored according to their tumor archetype in patients tumor sections. (**f)** Hierarchical clustering (quantized Wasserstein distance and Ward clustering) of the tumor archetypes identified with the T-UNI model using full retraining, KoLeo loss and 16K prototypes within each patient. Top: in the T-UNI embedding space. Bottom: in the raw gene expression space.

**Figure 5.**
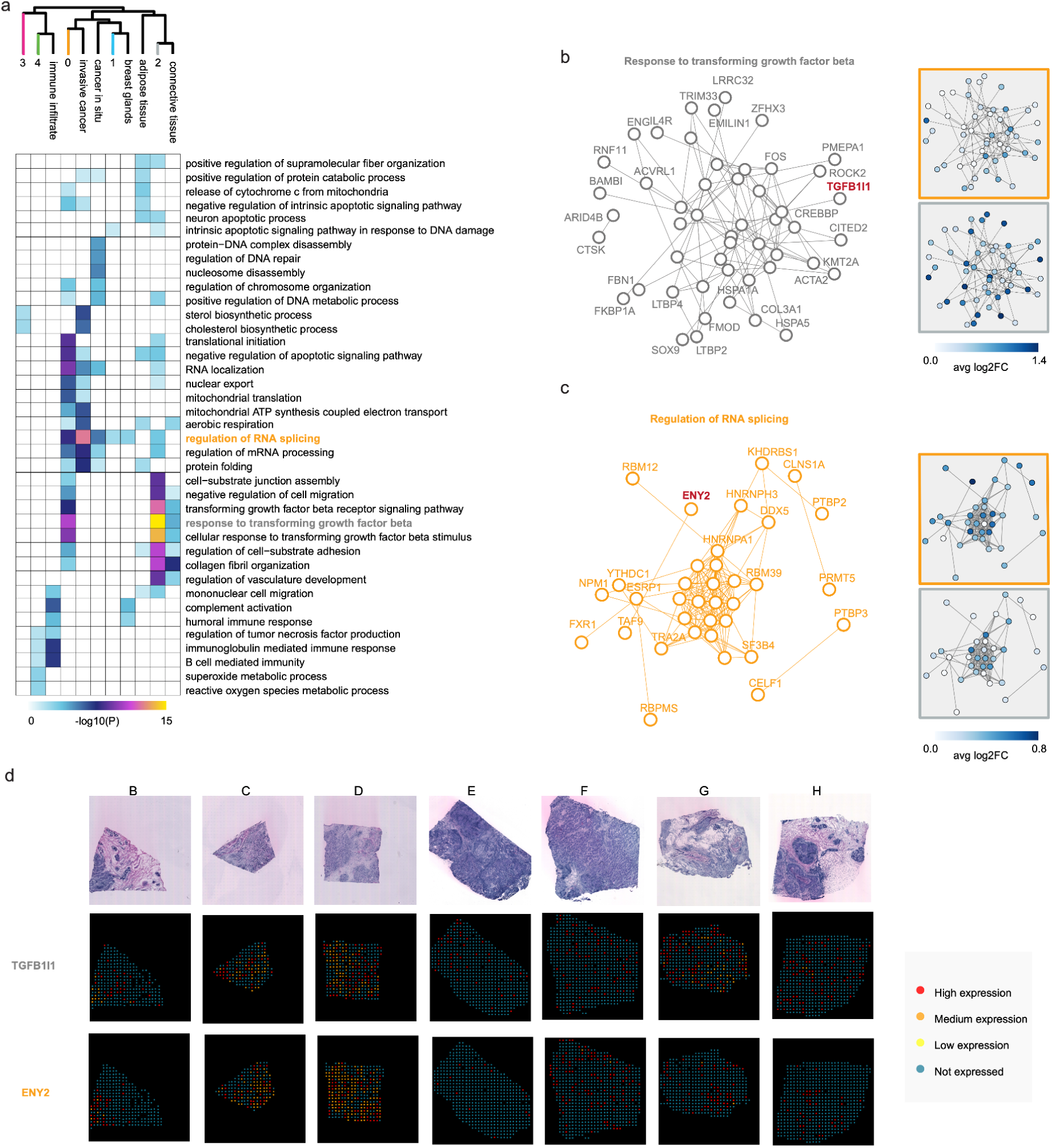
| Histopathology-derived tumor archetypes reveal distinct molecular pathways in invasive breast cancer. (**a**) Heatmap of the biological pathway enrichment analysis of differentially expressed genes (using MAST^66^; log2FC>0.25 and P-value<0.01) between the different tissue types annotated by pathologists as well as tumor archetypes identified with T-UNI full training strategy, KoLeo loss, 16K prototypes. The –log10 pvalue is reported for the top 10 significantly enriched pathways in each tumor archetype or tissue type, provided they contain at least 5 of the identified marker genes. Hierarchical clustering is performed using 1-pearson correlation as the distance metric, and the complete method from *hclust* function in the *stats*^74^ R package. (**b**) Network of protein-protein interactions for marker genes for tumor archetype 2, involved in the response to transforming growth factor beta pathway, together with their average log2 fold change in tumor archetype 0 and 2 compared to all the others. Edges represent experimentally determined protein-protein interactions annotated in the STRING database ^75^. Nodes indicate proteins. (**c**) Same as **b** for marker genes for tumor archetype 0 that are involved in the regulation of RNA splicing pathway, together with their average log2 fold change in tumor archetype 0 and 2 compared to all the others. (**d**) Spot expression of TGB1I1 and ENY2 involved in TGF-β and RNA splicing pathways respectively as shown in **b** and **c**. For each patient, the distribution of the given gene’s expression was analyzed in log2 scale. The mean and standard deviation (std) were calculated. The expression levels of each spots were then categorized as follows: Not expressed if *log*_2_ (*count* + 1) = 0; Low expression if *log*_2_ (*count* + 1) < *mean* − *std*; Medium expression if*mean* − *std* < *log* (*count* + 1) < *mean* + *std*; High expression if *log*_2_ (*count* + 1) > *mean* + *std*, with *count* being the raw count of the spot for the given gene.

Together, these results provide the first evidence that RNA metabolism and TGF-β signaling co-exist within invasive breast cancer tissue as distinct, well-defined regions, differing in abundance, spatial distribution, and morphological characteristics. Notably, these specialized tumor compartments are recoverable only by a foundation model (hFM) trained to specialize in this specific tissue type. This highlights the power of AI to discriminate adjacent tumor regions that are nearly indistinguishable morphologically yet encode relevant molecular differences, a capability that has not been previously achieved using conventional histopathological analysis.

## Discussion

In recent years, histopathology foundation models have rapidly evolved in size and complexity, demonstrating strong performance in tasks such as segmentation, cancer diagnosis and classification, treatment response, biomarker discovery, and outcome prognosis ^17,22,50–52^. However, critical gaps remain in understanding which biological concepts, spanning tissue, cellular, and molecular levels, are actually captured within their learned representations.

Our work aims at filling these gaps through comprehensive and original studies while focusing on breast cancers. First, we carried out a comprehensive benchmark of state-of-the-art hFMs across biological scales, enabling us to rank the aspects that are primarily encoded by these models. In particular, we have identified that knowledge of nuclei composition, overall texture and colour properties predominate. Notably, the high sensitivity of these models to color variations may help explain the strong patient-specific effects we observed. Since H&E staining is not standardized across laboratories, such technical variability is likely embedded in the learned representations. As a result, hFMs may inadvertently prioritize staining artifacts over biologically relevant features. This underscores the importance of accounting for color variability during model development. Indeed, our extended pre-training strategies help mitigate patient-specific effects by encouraging the model to focus on features that are consistent across patients, rather than spurious differences such as those arising from variations in staining concentration. Secondly, we proposed a semanticity measure based on the Shannon Entropy, enabling unsupervised model selection, which correlates significantly with the ability of models to discriminate the aforementioned features and tissue types. Nevertheless, we demonstrated that these models favour a global understanding of tissues to the detriment of the ability to encode distinct tumor regions with specific molecular profiles, a capability deemed crucial for characterizing ITH and improving patient stratification and therapy response prediction.

We therefore performed an in-depth analysis of several extended pre-training strategies, which differ in their ability to retain potentially relevant prior knowledge, to efficiently specialise models in encoding tumours. To this end, we identified that a global update of the model weights with KoLeo regularization was the best at optimizing a trade-off between tissue clustering, batch effects and encoding of handcrafted features. Newly learned representations were further used to uncover tumor archetypes which relate to molecularly distinct tumor tissues, indicating that molecular alterations manifest as morphological tissue patterns recognizable by our specialized models ^27–31^. In particular, we unveiled the molecular identities of the archetypes by analyzing their gene expression profiles using spatial transcriptomics (ST) data matched to the analyzed H&E tumor patches. Notably, we found that the most present archetype exhibits enrichment in aberrant mRNA metabolism, including RNA splicing/localization and dysregulated energy metabolism. Whereas the second most prevalent archetype is characterized by TGF-β signaling activation, a pathway critical to tumor progression, metabolic reprogramming, and the tumor microenvironment. While RNA splicing dysregulation and TGF-β signaling are both established hallmarks of cancer^43,46,53,54^, their spatial co-occurrence within defined tumor regions has not been systematically explored. Our findings thus provide novel insights into tumor heterogeneity, revealing distinct morphological domains associated with these pathways. Future studies will determine whether the relative abundance of these clusters correlates with specific clinical outcomes or serves as a predictor of therapy response.

We believe that our methodological contributions on both computational biology and machine learning can pave the way to many future works. For instance, we plan to extend our analysis to the single-cell level by studying cell representations derived from token ones, already present in hFMs, which could capture finer semantics than the patch representations considered in our study. In addition, although our current results are all the more attractive for having required few computational resources, we wish to significantly increase the number of patients studied in order to identify richer archetypes. To this end, it may also be relevant to consider multi-modal pathology foundation models, like UNI2-h that included immunohistochemistry (IHC) slides or potentially new ones incorporating multiplex data. Additionally, we acknowledge that accurately assessing cellular heterogeneity, a key driver of both tumorigenesis and drug resistance, alongside interactions within the tumor microenvironment is essential for refining patient stratification and developing effective therapies. Therefore we envision exploring the clinical implications of the computationally defined tumor archetypes, investigating their composition, association with patient outcomes, and therapy response. Finally we emphasize that these findings may be transformative for ST data analysis. Existing approaches for analyzing ST data rarely integrate information from histopathology. When they do, most methods use spatial organization to denoise molecular data^55,56^. However, spatial proximity alone is not always biologically meaningful in tumor biology. An alternative strategy relies on manual tissue region segmentation by pathologists, followed by differential expression analysis based on these expert-defined labels. However, this approach is limited by subjectivity and predefined tissue categories, potentially overlooking biologically relevant but visually subtle tumor subtypes. Here, we propose a novel method that enables the unsupervised labeling of molecular spots based on H&E image-derived features. This approach combines the strengths of both strategies, by aggregating spots to increase sensitivity in marker identification while assigning labels based on morphological appearance rather than predefined human annotations or marker gene selection. By leveraging histopathology-driven insights, this method enhances the biological interpretability of spatial transcriptomic data, providing a data-driven alternative to manual tissue annotation.

## Methods

### Datasets

#### Full dataset

We used a publicly available dataset, the HER2-positive breast cancer dataset^35^ composed of 36 whole slide images collected across 8 patients (resolution: 1μm/pixel). One image per patient has been annotated by a pathologist. Given the theoretical ST spot diameter of 100µm and the resolution of the HER2-positive breast cancer dataset of 1µm/pixel, the resulting squared spot patches are 100 pixels on each side. Therefore, patches of size 100×100 pixels without overlap have been extracted from the images to match the spatial transcriptomic spots size, representing a total of 103’023 patches. Patient A has been excluded for suspicion of technical bias in the acquired data (**Fig S1a**). Among the 103’023 patches, 11’582 are spots with associated RNA-sequencing data, of which 3’139 are labeled.

#### Invasive cancer patches dataset

To further explore intra-tumor heterogeneity, invasive cancer patches have been extracted from the full dataset, first considering non-overlapping 100*100 tiles within corresponding regions. To augment our subset of invasive cancer patches and potentially overcome imprecision of region annotations at the patch level, we applied the k-nearest neighbors (*k*-NN) algorithm to all patches in the full dataset using their labels for training, while benchmarking patch representations from 6 hFMs. All patches then classified as invasive cancer were taken as the effective subset for our analysis.. The chosen optimal *k* for *k*-NN corresponds to the one maximizing the median F1 score across patient after testing different values of *k* ranging from 3 to 21. While the median F1 scores are similar across the different hFMs (**Fig S6b**), we utilized predictions resulting from SimCLR representations (median F1 score of 0.82; median accuracy of 0.84). The final invasive cancer patches dataset contains 54,311 patches, including 5,683 spots among which 1,571 are labeled and 143 (9%) have been re-annotated by an expert pathologist. This invasive cancer patches dataset was further used to retrain the T-UNI models.

### Shannon entropy of explained variance

To evaluate the distribution of variance among the SVD singular vectors of the precomputed image representations, the normalized shannon entropy *H* of the explained variance vector is computed as follows:

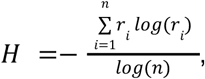

where *r_i_* represents the proportion of the total variance explained by the i-th singular vector, and *n* is the total number of singular vectors. A low Shannon entropy indicates that a few components account for most of the variance in the data. In contrast, a high Shannon entropy suggests that the variance is spread across multiple dimensions.

### Handcrafted feature extraction

Segmentation of the nuclei has been performed using CellViT^25^. Features related to the morphology and texture of the nuclei have been extracted using scMTOP^57^ and correspond to the “Nuclei-Morph”, and “Nuclei-Texture” features respectively. We have enhanced the “Nuclei-Morph” category by computing Zernike moments of each cell using the Mahotas python package^58^. We have adapted scMTOP to add color descriptors of the nuclei, referred to as “Nuclei-Color” features, namely mean, skewness, kurtosis and entropy of each RGB color channel, as well as intensity and transparency. Finally, statistics on the cell type composition have been computed based on cell labels outputted by CellViT, and are referred to as “Nuclei-Composition”. Features related to the extracellular matrix color (“ExtraCell-Color” features) and texture (“ExtraCell-Texture”) have been computed using scikit-image^59^. Similarly, texture and color features at the entire patch level have been extracted and are referred to as “WholePatch-Texture” and “WholePatch-Color”. The list of all handcrafted features and their descriptions is available in **Supplementary Table2**.

### Regression for handcrafted features prediction

To elucidate a part of the semantic information encoded by hFMs, we predicted each extracted handcrafted feature individually from hFM representations using linear regressions. We employed a 5-fold cross-validation approach and reported the mean R² across the five folds for each handcrafted feature.

### Clustering of different tissues

Clustering was performed on the image representations obtained from various hFMs or from our selected handcrafted features taken as a whole. Performance was measured using Adjusted Rand Index (ARI) scores computed using the original tissue types as true labels (“*ARI with tissue type*”). Before applying the k-means clustering algorithm with k set as the known number of classes, 2-dimensional UMAP embeddings were computed to account for non-linear relations between image representations, validating various hyperparameters: *n_neighbors* ranging from 10 to 400, and *min_dist* values of 0.001 and 0.1. For each configuration, hyperparameters maximizing the *ARI with tissue type* score were selected. Clustering is performed either on the whole dataset or for each patient individually. The UMAP-k-means approach was compared to raw-k-means where k-means is directly computed on the raw image representations, and to SVD5-kmeans which factor image representations using the first 5 principal components of SVD before performing k-means. UMAP-kmeans was chosen for its superior performance (**Fig S2d, Supplementary Table4**).

### Extended pre-training of the UNI foundation model

Extended pre-training of UNI has been performed only on our selected invasive cancer patches to refine their learned representations. Two different methods have been employed and compared: 1) ExPLoRA^39^, a state-of-the-art technique based on LoRa^60^ which performs extended pre-training updating only a small fraction of the initial foundation model weights; 2) the standard full extended pre-training of the model, where all weights are updated. For each strategy, we compared two different regularization losses (KoLeo and KDE), as well as the number of prototypes these models depend on. The list of all models is provided in **Supplementary Table3**.

#### Regularization losses

DINOv2^61^, the self-supervised learning approach utilized by UNI^44^, employs the Kozachenko–Leonenko (KoLeo)^41,42^ regularization loss, which aims at maximizing differential entropy via a convex regularizer applied independently over each batch sampled during training. It is defined as follows:

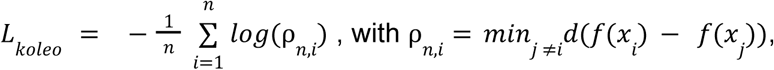

with *n* the number of samples, *f*(*x_i_*) the hFM representation of the *i*-th data point and *d* is the cosine distance.

By maximizing the distances between neighboring vectors, the KoLeo loss ensures that the learned representations are uniformly distributed within their space. However, Zimmermann et al.^34^ argue that while KoLeo has been demonstrated to be effective in natural image pretraining with high sample diversity, when samples get very similar, such as in digital histopathology, the loss can approach infinity. Therefore, they suggest using the kernel density estimator (KDE) with the unnormalized von Mises-Fisher (vMF) kernel, which allows similar images to cluster more smoothly in the embedding space depending on the choice of inner kernel.. The KDE loss is defined as follows:

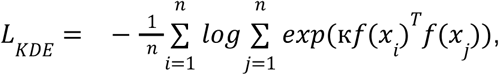

with к a scaling constant and other terms are defined above. Both the KoLeo and KDE losses are tested in our analysis.

#### Prototypes

DINOv2 depends on the online clustering scheme SwAV^62^, which relies on learned prototypes, estimating centroids on the fly within the representation space learned by the model. These prototypes promote the formation of clearly distinct regions, each encoding distinct high-level concepts or patterns identified by the model. Therefore, the number of prototypes influences the level of fine-grained-ness of the clusters. In theory, a greater diversity of training images necessitates a larger number of prototypes to accurately cluster these images. However, evidence suggests that an excessive number of prototypes may lead to significant redundancies among them, a phenomenon termed as *partial prototype collapse*^63^. Additionally, increasing the number of prototypes incurs significantly higher computational costs. While UNI has been trained using 65536 prototypes, we decided to perform the extended pre-training using various prototype numbers: 4096, 16384 and 65536.

#### Hyperparameters selection

Apart from the different losses tested and the various prototype numbers, the same parameters as those chosen in UNI^17^ (available on their <u>github repository</u>) were initially adopted, with a few modifications implemented. First, the warmup phase, typically used to gradually increase the learning rate at the beginning of training, was omitted because UNI had already undergone pretraining. The cosine decay schedule was started at a learning rate of 1 × 10^-4^. Secondly, the final layer of the prototype was frozen after a significant increase in training loss was observed during its training (**Fig S6a**). Finally, the number of epochs was restricted to one, as no further improvement in the total training loss was detected beyond this point and the batch size was set to 40 to fit within memory constraints. For all models, the extended pre-training lasted less than one hour on our commercial RTX 3090 GPU.

### Identification of invasive cancer archetypes

Clustering was performed with the UMAP-k-means strategy on a newly defined subset of invasive cancer patches to identify tumor archetypes. In particular, we observed that models resulting from extended pre-training (especially full training) on the first invasive cancer subset become significantly better at distinguishing them from the other tissue types. Hence we used these refined models to get a better curated subset of invasive cancer patches following the same k-NN strategy than defined above. Then, hFM representations were embedded using UMAP embeddings, testing various UMAP hyperparameters similarly than for previous experiments. However, hyperparameter validation for both UMAP and k-means was performed differently, toward maximizing jointly 1) the silhouette score, and 2) the “*batch effect mitigation*” score, defined as 1-”*ARI with patient*”, computed using patient labels instead of tissue labels on patches from invasive cancer. A high batch effect mitigation score indicates low batch effect. To simultaneously optimize both objectives, we calculated the Euclidean distance from the obtained silhouette scores and batch effect mitigation scores to the optimal point (1, 1). This point represents the situation of perfectly defined clusters (silhouette score = 1) that are also perfectly balanced across patients (batch effect mitigation score = 1) (**Supplementary Fig.4a**).

### Molecular characterization of the image tumor archetypes

#### Filtering

The processed gene count matrices deposited on Zenodo were used^35^ (https://zenodo.org/records/4751624). Spots with no reads were excluded. For each patient, a pseudo-bulk expression vector was created by aggregating the gene expression vectors from all spots. These vectors were then log2-transformed. The resulting gene read density showed a bimodal distribution: a low-density peak for low-abundance genes or those with spurious mappings, and a high-density peak for reliably expressed genes. To identify these reliably expressed genes in each patient, a two-component Gaussian mixture model was applied to the log2-transformed pseudo-bulk expression vector using the R package mclust^64^ (**Supplementary Fig.5b**). A gene was kept if in all 7 patients, it has less than 1% chance of belonging to the low-density peak, resulting in a final set of 11,591 genes and 11,554 spots. The spatial transcriptomics (ST) data was then treated as single-cell data and the *NormalizeData* function from Seurat^65^ was used to normalize the data: gene counts for each spot were divided by the total counts of that spot and multiplied by 10,000, followed by log transformation. The resulting filtered expression matrix is referred to as the “*gene expression space*”.

#### Uncovering tumor archetype-specific biological pathways

In order to characterize molecularly the tumor archetypes identified by the vision encoders, we extracted the tumor spots and performed a differential gene expression (DGE) analysis between tumor archetypes using the *FindAllMarkers* function from Seurat^65^, with patients specified as latent variables, to identify differentially expressed genes while mitigating the patient batch effect present in the filtered gene expression space (**Fig S4c**, *upper*). The MAST^66^ method was used for differential gene expression (DGE) analysis. Up-regulated marker genes for each tumor archetype have been extracted by selecting the genes expressed at least in 25% of the spots of the archetypes tested, with a minimum log2 fold-change of 0.25 and a p-value of 0.01. Finally, using g:Profiler^67^, we extracted for each tumor archetype the top 10 significantly enriched GO biological pathways^68^ containing at least 5 of the identified marker genes. Redundant pathways have been curated manually. The exact same analysis has been performed to characterize the annotated tissues.

#### Towards a batch-corrected gene expression space

To obtain a batch-corrected gene expression space, we utilized the ComBat^69^ method from the sva^70^ package, originally developed for microarray data. Although various batch correction methods were tested, ComBat demonstrated the most effective batch removal. Applying ComBat to the gene expression matrix resulted in a matrix with the same dimensions (11,591 genes and 11,554 spots), which we refer to as the “combat-corrected gene expression space” (**Supplementary Fig.4c**, *lowe*r).

### Quantized Wasserstein distance

We use the Wasserstein distance, also known as the Earth Mover’s Distance, to assess the dissimilarity between tumor archetypes, i.e the singular tumor clusters identified by the previously defined strategies, in both image or molecular representation spaces. This distance, based on Optimal Transport, measures the minimal total distances to be covered to transport each point of a distribution to those of another^71^. Its computation is cubic in the number of samples when both distributions have similar sizes, hence given the high number of samples identified in our tumor archetypes, we estimated it using the *quantized* Wasserstein distance. It comes down to first reducing each distribution, by taking their k-means centroids as support and assigning them a relative importance which coincides with cluster proportions. Then factored distributions were compared using their Wasserstein distances, leveraging the implementation of the Python Optimal Transport library^72^. To improve the estimation of the true Wasserstein distances, a high number of 5000 clusters for each distribution was taken.

Remark that different hFM representation spaces can have different volumes but similar discernibility between archetypes or clusters (**Fig. 4a**). Hence for comparing such spaces in an unbiased manner, the quantized Wasserstein distance was computed on normalized representations. Specifically, we computed the maximal diameter α of the UNI representation space by finding the maximal distance between the two most distant points, and normalized the other embedding spaces by multiplying them by their maximal diameters and dividing by α.

### Hierarchical clustering

Hierarchical clustering between tumor archetypes is computed in the image or gene expression embeddings. In both cases, it is computed using the ward method and the quantized Wasserstein distance, using the *linkage* method from scipy^73^.

### Data availability

The data and the model weights of T-UNI are available via Zenodo (https://zenodo.org/records/15053890). The processed count matrices derived from the raw ST data and the associated brightfield images (HE-images) were downloaded from Zenodo (https://doi.org/10.5281/zenodo.4751624) ^35^.

### Data availability

The source code is available via Github (https://github.com/idiap/TumorArchetype-FM).

## Acknowledgements

This work was financially supported by Novartis Foundation for Biomedical Research grant no. 22B104 and by Idiap Research Institute (KPI-Boost) (to LF); The Personalized Health and Related Technologies PHRT Project 2022-644 (to CVC); Idiap research institute (to VJ). Figure 1 was created with BioRender.com. We thank Mateo Thonet for integrating UNI and Prov-GigaPath into the code.

## Ethics declarations

The authors declare no competing interests.

## FIGURES AND TABLES

**Supplementary Figure 1.**
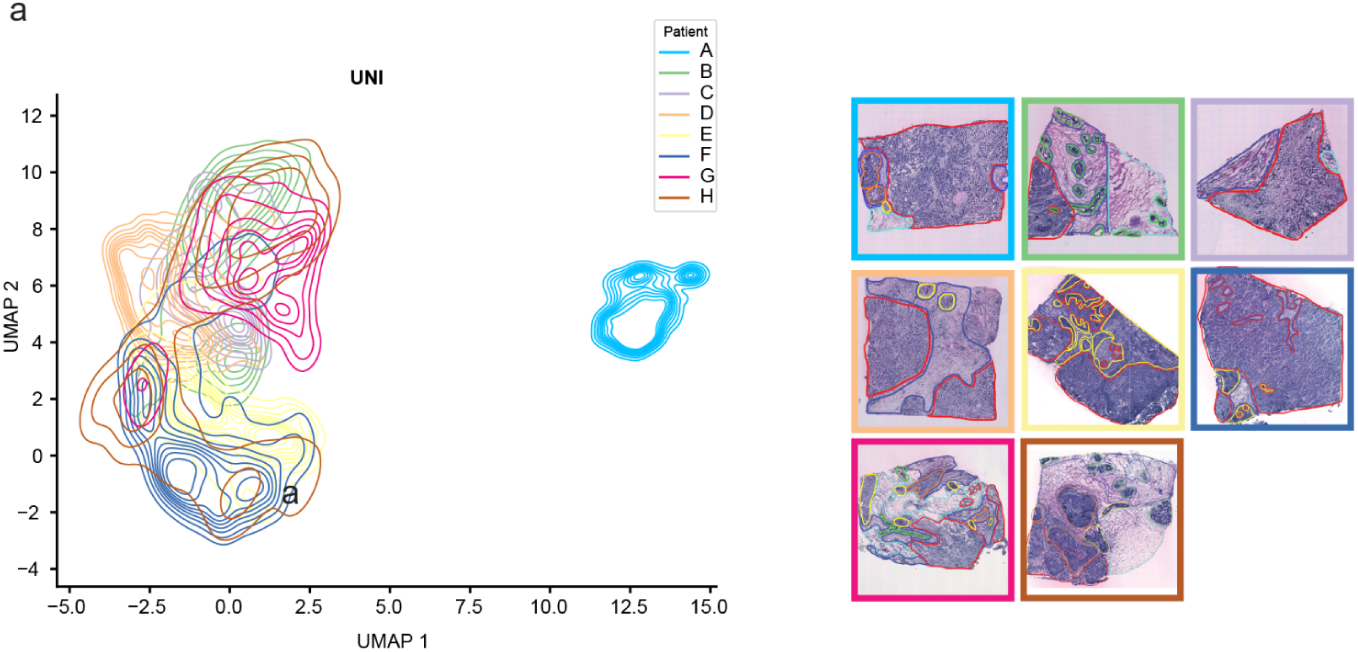
| **a** UMAP projections^36^ of all patches (*n*_*patc*ℎ*es*_ = 103023) embedding extracted from UNI, showing patient A as an outlier in the data.

**Supplementary Figure 2.**
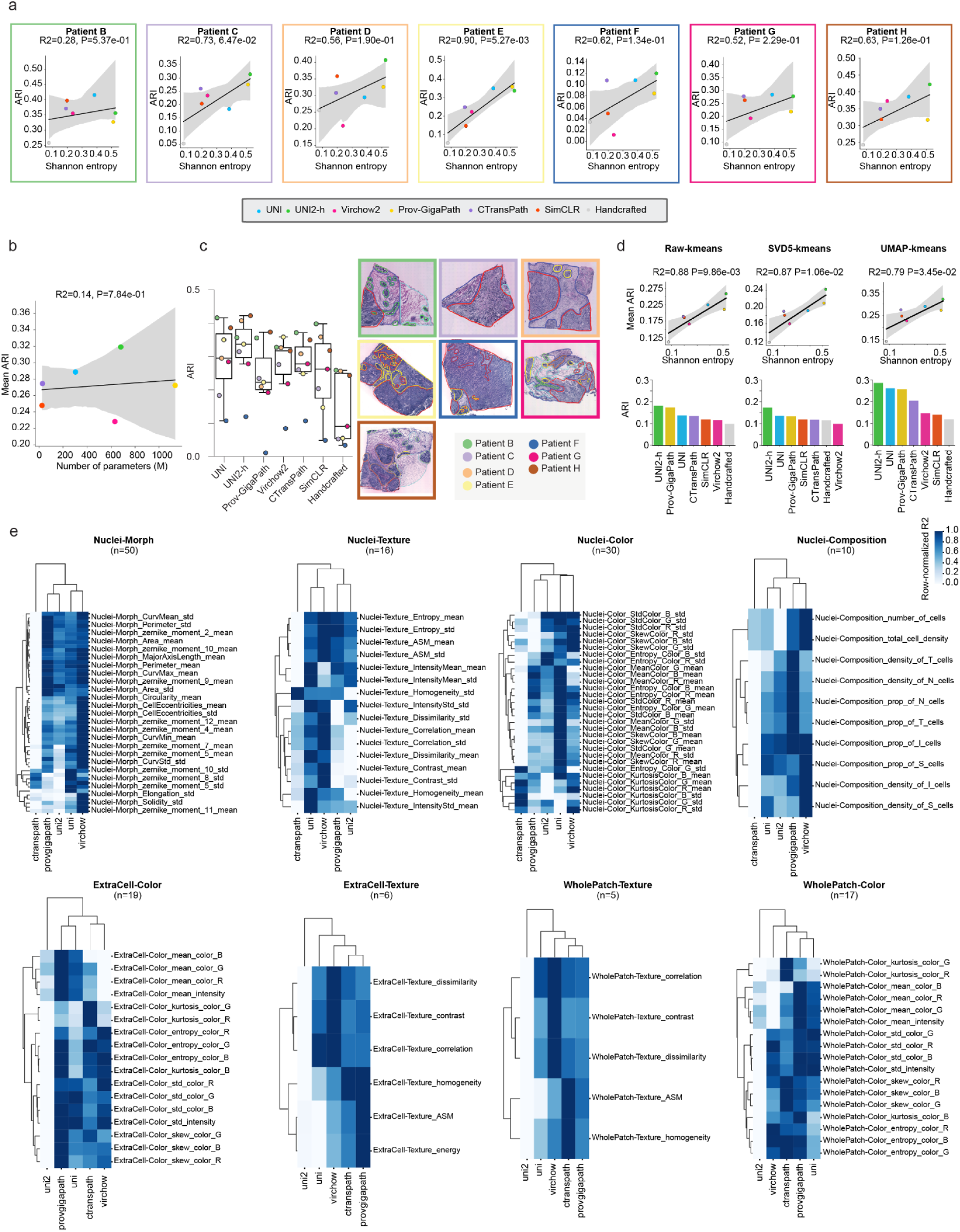
| **a**. Correlation between SE and ARI score for each patient for the different hFMs and handcrafted features. ARI measures tissue type discrimination. R² indicates the Pearson correlation coefficient. The line marks the linear regression fit with the 95% confidence interval as error band. Clustering technique used is UMAP k-means. **b**. Pearson correlation between the number of parameters of the hFMs and the mean ARI score across patients. ARI measures tissue type discrimination. Clustering technique used is UMAP k-means. **c**. ARI scores measuring tissue type discrimination per patient for the different hFMs and handcrafted features, together with the whole slide image to illustrate the clustering task. Clustering technique used is UMAP k-means. **d**. Top: Correlation between SE and mean ARI score across patients. Bottom: Overall ARI score on all patches, measuring tissue type discrimination for different methods. Left: k-means is applied on the raw embeddings extracted from the hFMs or handcrafted features. Middle: k-means is applied on the first 5 principal components from singular value decomposition of the raw hFMs embeddings or handcrafted features. Right: UMAP has been applied to the raw hFMs embeddings and handcrafted features prior to applying k-means. The reported ARI scores are the best scores obtained across all UMAP parameters tested, with *n_neighbors* ranging from 10 to 400, and *min_dist* values of 0.001 and 0.1 (see *Materials and Methods* section). **e**. Heatmaps of R² score from linear regression predictions of handcrafted features based on the different hFMs patches embeddings, grouped by feature category (nuclei morphology, nuclei texture, nuclei color, nuclei composition, extracellular color, extracellular texture, color and texture of the whole patch). The R² scores have been normalized over rows to compare models between them. SimCLR has been excluded from the heatmaps since displaying systematically lower R².

**Supplementary Figure 3.**
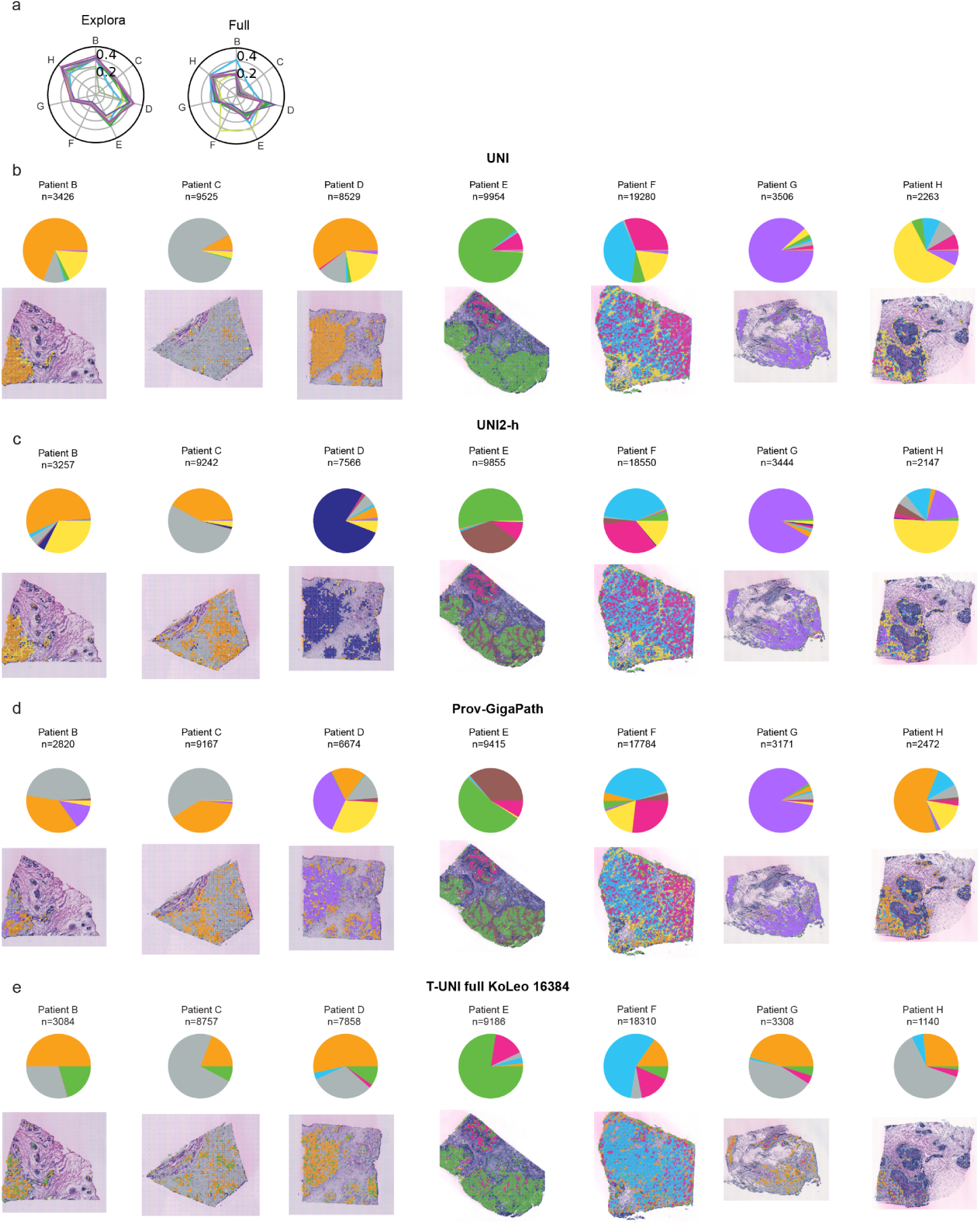
| **(a)** Radar plot showing ARI scores per patient measuring capacity of UNI and T-UNI models to discriminate tissue types. Clustering technique used is UMAP k-means. **(b)** Pie chart showing the relative abundance of the 7 tumor archetypes identified with UNI. Top: Relative abundance of morphological tumor archetypes per patient. Bottom: Invasive cancer patches colored according to their tumor archetype in patients tumor sections. **(c)** Same as b but for the 9 tumor archetypes identified with UNI2-h. **(d)** Same as b but for the 8 tumor archetypes identified with Prov-GigaPath. **(e)** Same as b but for the 5 tumor archetypes identified with our T-UNI, full retraining, KoLeo regularization and 16384 prototypes.

**Supplementary Figure 4.**
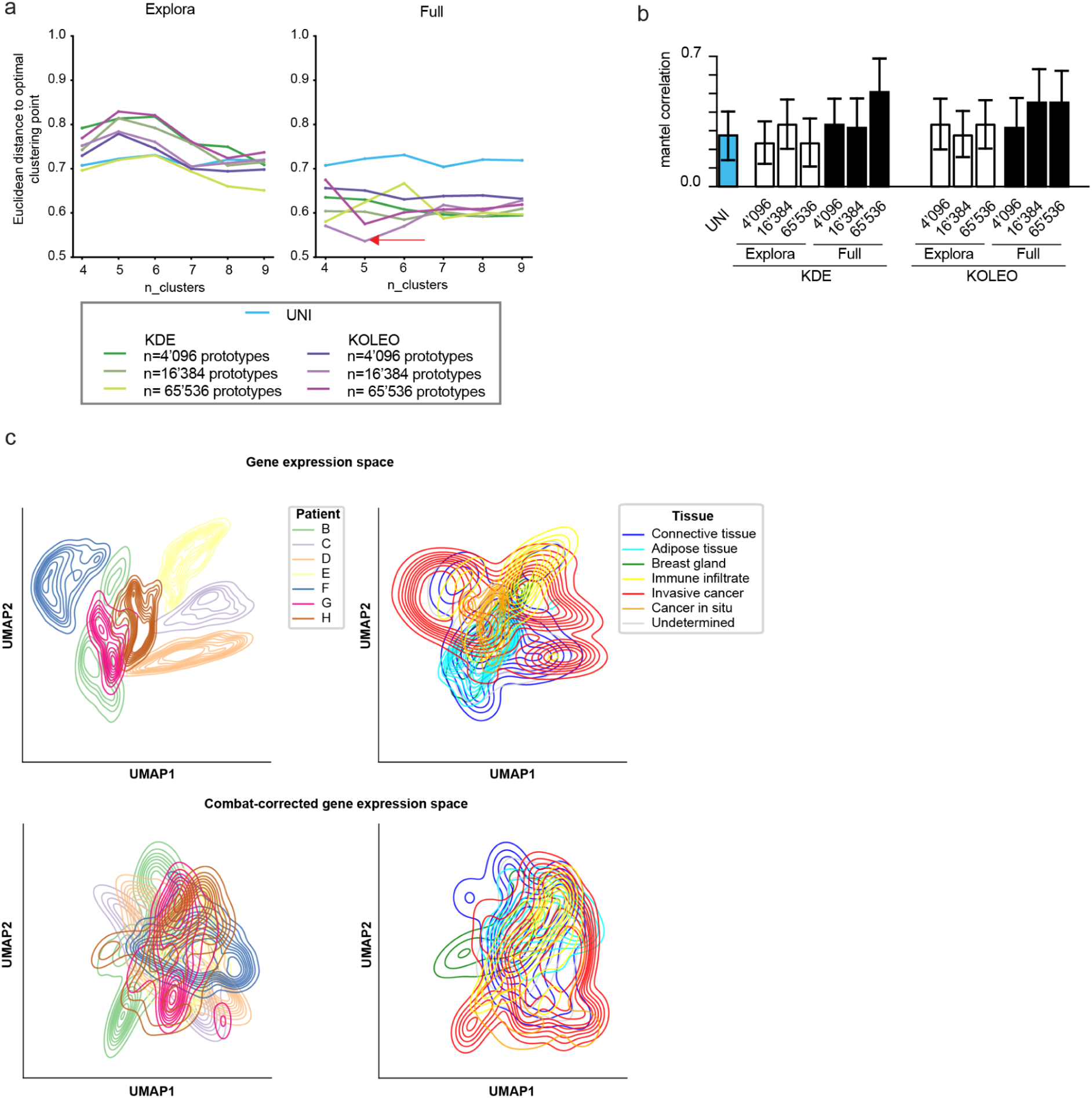
| **a**. Euclidean distance for each value of k between the point defined by the silhouette score and batch effect mitigation score, and the optimal point (1, 1). This point represents the situation of perfectly defined clusters (silhouette score = 1) that are also perfectly balanced across patients (batch effect mitigation score = 1). Clustering is performed using UMAP-kmeans, selecting the UMAP parameters leading to the best silhouette score for each k for T-UNI models trained using ExPloRa (left) and using full retraining (right). The red arrow shows the model and number of clusters being the closest to the optimal clustering. **b**. Bar graphs of average mantel correlations between molecular hierarchies across patients for UNI and the T-UNI models. Molecular hierarchies are computed in the gene expression space using quantized Wasserstein distances. 95% confidence intervals are reported in black. **c**. Kernel density plots of UMAP projections of the filtered gene expression space, composed of 11,591 genes and 11,554 spots (upper) and its ComBat^69^-corrected version (lower). Left: colored by patient origin. Right: colored by tissue origin.

**Supplementary Figure 5.**
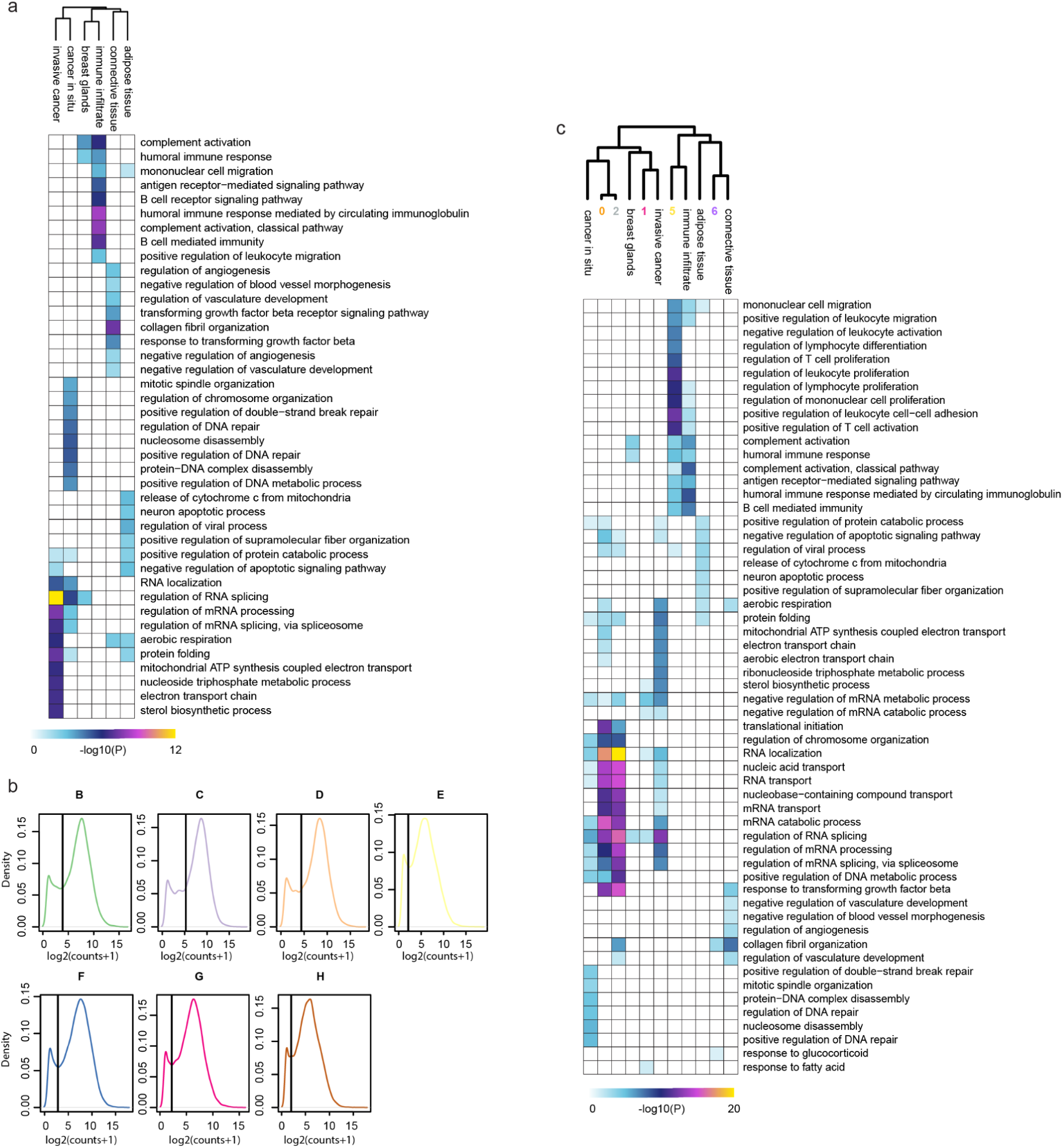
| **a**. Heatmap of the biological pathway enrichment analysis of differentially expressed genes (using MAST^66^; log2FC>0.25 and P-value<0.01) between the different tissue types annotated by pathologists. The –log10 pvalue is reported for the top 10 significantly enriched pathways in each tissue type, provided they contain at least 5 of the identified marker genes. Hierarchical clustering is performed using 1-pearson correlation as the distance metric, and the complete method from *hclust* function in the *stats*^74^ R package. **b**. Log2-transformed gene read density per patient, showing a bimodal distribution: a low-density peak for low-abundance genes or those with spurious mappings, and a high-density peak for reliably expressed genes. To identify these reliably expressed genes in each patient, a two-component Gaussian mixture model was applied to the log2-transformed pseudo-bulk expression vector using the R package mclust^64^. A gene is considered as reliably expressed if it has less than 1% chance of belonging to the low peak (threshold indicated by the black vertical line). **c**. Heatmap of the biological pathway enrichment analysis of differentially expressed genes (using MAST^66^; log2FC>0.25 and P-value<0.01) between the different tissue types annotated by pathologists as well as tumor archetypes identified with UNI (baseline). The –log10 pvalue is reported for the top 10 significantly enriched pathways in each tumor archetype or tissue type, provided they contain at least 5 of the identified marker genes. Hierarchical clustering is performed using 1-pearson correlation as the distance metric, and the complete method from *hclust* function in the *stats*^74^ R package.

**Supplementary Figure 6.**
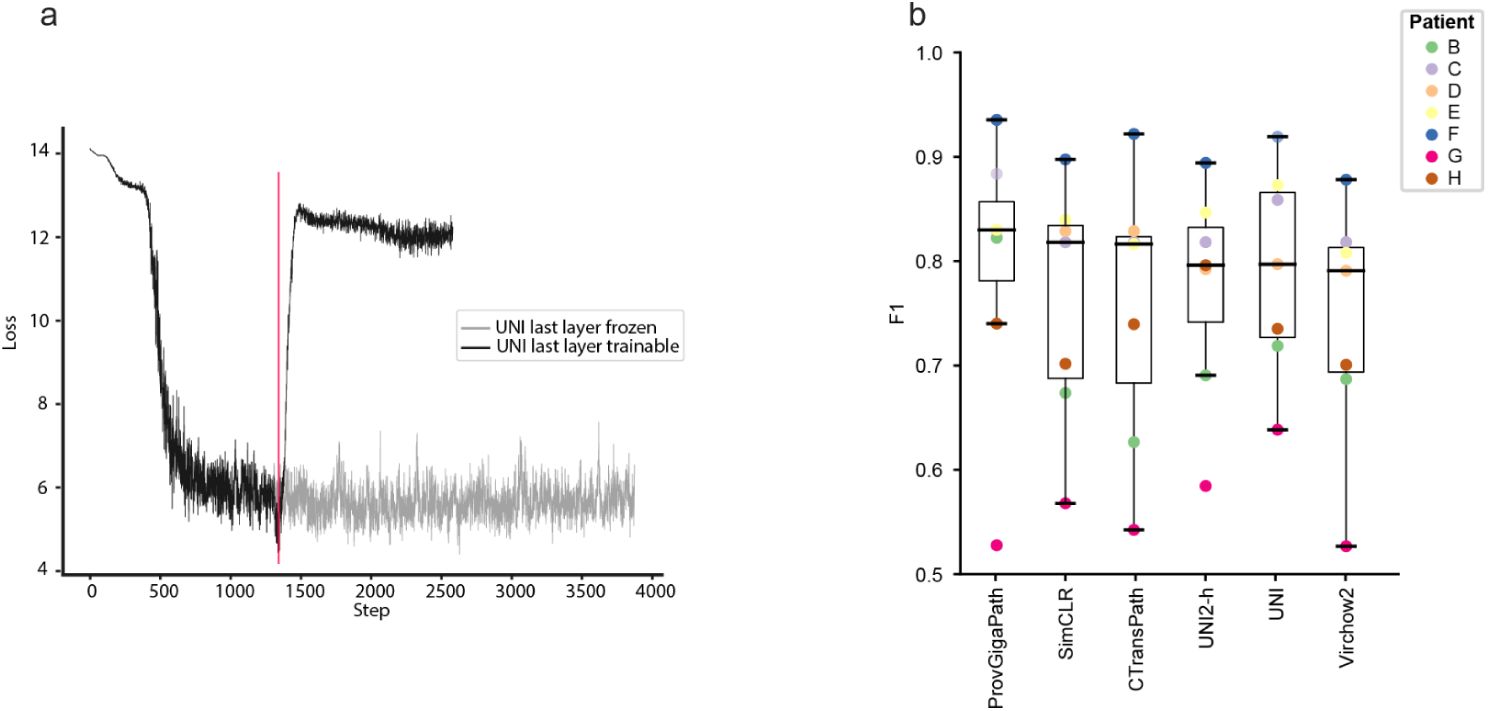
| **a**. DinoV2 total loss when performing extended pre-training with invasive cancer patches. Two different set-ups are shown: the last layer is made trainable after one epoch (black) and the last layer remains frozen (gray). The vertical red line indicates the first epoch. **b**. F1-score per patient after k-NN relabeling of patches per patient, performed on the different hFMs.

## SUPPLEMENTARY TABLES LEGENDS

**Supplementary Table 1** Histopathology foundation models (hFMs) descriptions (benchmark tasks, training data, model architecture) together with their performances on the HER2 dataset (Shannon Entropy and ARI scores).

**Supplementary Table 2** List and description of handcrafted features computed after CellViT^25^ segmentation.

**Supplementary Table 3** List and parameters of the different extended-trained UNI (T-UNI) models. Extended pre-training has been performed using the same invasive cancer patches dataset for all models (see *Materials and Methods* section)

**Supplementary Table 4** Model performances on the HER2 dataset^35^ for the 6 baseline hFMs and the 12 T-UNI models.

**Supplementary Table 5** Description of the patch dataset created from the HER2 dataset^35^, with number of patches and labeled spots per tissue type and per patient.

